# Chemobiosis reveals tardigrade tun formation is dependent on reversible cysteine oxidation

**DOI:** 10.1101/2023.05.15.540801

**Authors:** Amanda L. Smythers, Kara M. Joseph, Hayden M. O’Dell, Trace A. Clark, Jessica R. Crislip, Brendin B. Flinn, Meredith H. Daughtridge, Evan R. Stair, Saher N. Mubarek, Hailey C. Lewis, Derrick R.J. Kolling, Leslie M. Hicks

## Abstract

Tardigrades, commonly known as ‘waterbears’, are eight-legged microscopic invertebrates renowned for their ability to withstand extreme stressors, including high osmotic pressure, freezing temperatures, and complete desiccation. Limb retraction and expelling their internal water stores results in the tun state, greatly increasing their ability to survive. Emergence from the tun state and/or activity regain follows stress removal, where resumption of life cycle occurs as if stasis never occurred. However, the mechanism(s) through which tardigrades initiate tun formation is yet to be uncovered. Herein, we use chemobiosis to demonstrate that tardigrade tun formation is mediated by reactive oxygen species (ROS). We further reveal that tuns are dependent on reversible cysteine oxidation, and that this reversible cysteine oxidation is facilitated by the release of intracellular reactive oxygen species (ROS). We provide the first empirical evidence of chemobiosis and map the initiation and survival of tardigrades via osmobiosis, chemobiosis, and cryobiosis. *In vivo* electron paramagnetic spectrometry suggests an intracellular release of reactive oxygen species following stress induction; when this release is quenched through the application of exogenous antioxidants, the tardigrades can no longer survive osmotic stress. Together, this work suggests a conserved dependence of reversible cysteine oxidation across distinct tardigrade cryptobioses.

## Introduction

Tardigrades are a phylum of eight-legged microscopic invertebrates renowned for their remarkable ability to survive extreme environmental stressors.^1-6^ This survival is rooted in their ability to initiate cryptobiosis, a physiological state wherein metabolism slows to near undetectable conditions, enabling long-term survival despite inhospitable conditions.^7^ Although some eukaryotes and bacteria are capable of cryptobiosis, no eukaryotes are able to do so across the entirety of their lifespan, including as eggs, juveniles, and adults, or in response to such a broad range of stressors as tardigrades.^8^ Thus tardigrades’ ability to survive desiccation, freezing, oxygen starvation, fluctuating osmotic pressure, and ionizing radiation (via anhydrobiosis, non tun-forming cryobiosis, anoxybiosis, osmobiosis, and irradiation-induced dormancy, respectively) is matched by none.^9^ in response to such a broad range of stressors as tardigrades.^8^ Thus tardigrades’ ability to survive desiccation, fluctuating osmotic pressure, freezing, oxygen starvation, and ionizing radiation (via tun-forming anhydrobiosis and osmobiosis, and non tun-forming cryobiosis, anoxybiosis, and irradiation-induced dormancy, respectively) is matched by none.^9^

Mechanisms of tardigrade survival are still poorly understood. While several cryptobiotes rely on trehalose synthesis as a protectant during dormancy, many tardigrade species produce low or undetectable levels of the disaccharide.^10-14^ Tardigrades also lack highly conserved networks for stress regulation, including those that connect hypoxia, genotoxic stress, and oxidative stress to the conserved master regulator target of rapamycin (TOR).^15^ Instead, tardigrades rely on several organism-specific proteins for stress protection, such as the damage suppressor protein (Dsup) that associates with nuclear DNA to protect from ionizing radiation.^15^ Additionally, cytoplasmic-, secreted-, and mitochondrial-abundant heat soluble (CAHS, SAHS, and MAHS) proteins, collectively known as tardigrade disordered proteins (TDPs), possess sequences without conservation in other phyla and are essential for tardigrade survival to desiccation via anhydrobiosis.^16^ Further, late embryogenesis abundant (LEA) proteins and heat shock proteins (HSP) are widely distributed throughout tardigrade cells, and many are upregulated during tardigrade anhydrobiosis suggesting an essential role in cryptobiosis.^17-20^

The hallmark of most tardigrades undergoing cryptobiosis is their remarkable ability to shift into a shriveled anatomical state known as a tun, which they achieve by contracting their limbs, retract along their medial axis, and in the process expel their internal water stores.^21^ Upon exposure to favorable conditions, tardigrades rapidly distend and return to active metabolism. It has long been understood that tardigrades facilitate these anatomical transitions via active processes, yet the mechanism(s) through which tardigrades recognize environmental fluctuations and signal the transitions to enter and exit tuns remains largely unexplored.^22^ Tardigrades rely on mitochondrial activity for desiccation survival; using chemical inhibitors to uncouple the mitochondrial electron transport chain prevents tuns from forming following exposure to anhydrobiotic conditions.^23^ Further, the ATP-dependent differential phosphorylation of the AMP-activated protein kinase (AMPK) regulatory network following anhydrobiosis revealed that protein phosphatase 2A activation is essential to successfully induce tuns via anhydrobiosis.^24-26^ Both mitochondrial electron transport and AMPK regulation, both of which are highly conserved across taxa, engage in crosstalk with reactive oxygen species (ROS). In fact, mitochondrial ROS generation has been directly linked with AMPK activation in mouse embryonic fibroblasts, which is considered to be conserved across taxa.^27-30^ When combined with increased accumulation of antioxidant enzymes and the redox homeostasis mediator glutathione following anhydrobiosis,^31, 32^ evidence suggests oxidative signaling may play a key regulatory role in initiating tun formation.

Herein, novel evidence supporting reversible oxidation as an essential regulatory signal for cryptobiosis is demonstrated in the model tardigrade species, *Hypsibius exemplaris*. Tardigrades are shown to respond to exogenously applied ROS by forming tuns in a dose-dependent manner that is inhibited when cysteine thiols are irreversibly blocked. Thiol-specific fluorescent labeling enabled imaging via confocal fluorescence microscopy with quantification of reduced thiols accomplished through quantitative fluorescent assays. Rapid exposure to reducing conditions causes tun release and death, indicating that the oxidation and reduction of cysteine thiols is accomplished via highly regulated internal networks. Further, quantitative electron paramagnetic resonance spectrometry (EPR) spectroscopy reveals an intracellular release of ROS following stressor induction, that is fatal when blocked with exogenously applied antioxidants. Inhibition of voltage-dependent anion channel protein 2 generates tuns, suggesting ROS control of this ion transporter is likely implicated in tardigrade stress. Together, these data support reversible oxidative signaling as an indispensable regulatory mechanism for tardigrade survival to adverse environments, with a conserved role across distinct cryptobiosis.

## Materials and Methods

### Tardigrade husbandry

Cultures of *Hypsibius exemplaris* (Sciento; Manchester, UK) were reared in 1- or 2-L Erlenmeyer flasks on stationary-phase *Chlorella vulgaris*. Culture medium was changed biweekly using a 40-μm Corning^™^ Sterile Cell Strainers (Thermo Fisher Scientific, Waltham, MA), which retains the tardigrades while allowing algae and waste to flow through. The filter is inverted and rinsed into the same flask with the addition of fresh deionized water. Tardigrades are fed weekly with *C. vulgaris* that was grown photoautotrophically under previously established protocols.^23^ Cultures were maintained at room temperature on a 12:12 h light:dark cycle using a 7 W (630 lumens) LED lamp.^33^

### Cryptobiosis induction

Exposure time and concentration of each stressor was optimized to establish the highest post-cryptobiotic survival rates while maintaining high rates of cryptobiotic state induction. Survival for each stressor was defined as tardigrades exhibiting controlled movement of limbs and/or body post-recovery treatment. Each stressor was evaluated in four biological replicates of approximately 30 animals per replicate.

#### Cryobiosis

Tardigrades were transferred in droplets of minimal media (<40 μL) on a 35-mm plastic petri dish and placed in Thermo Fisher Scientific Nalgene ‘Mr. Frosty’ Freezing Container containing 70% isopropyl alcohol (Thermo Fisher Scientific, Waltham, MA) at -80 °C for 4 h. Tardigrades were subsequently removed from the apparatus, thawed at room temp, and monitored for recovery.

#### Osmobiosis (CaCl_2_ and sucrose)

Tardigrades were transferred in minimal media (<100 μL) on 35-mm plastic petri dishes and dosed with working concentrations of 75 mM CaCl_2_ (Amresco, Dallas, TX) or 600 mM sucrose (Millipore, Burlington, MA) for 5 and 1 h respectively and tun formation was observed. Animals were transferred to new plates containing culturing media and monitored for recovery.

#### Chemobiosis

Tardigrades were transferred in minimal media to 35-mm plastic petri dishes and dosed with working concentration of 750 μM hydrogen peroxide (Thermo Fischer Scientific, Waltham, MA) for 12 h and tun formation was observed. Animals were transferred to new plates containing culturing media and monitored for recovery.

### Tun blocking

*H. exemplaris* (30 animals per sample, n=4) collected in microcentrifuge tubes were subjected to a 30-min incubation in 30 μM iodoacetamide (IAM) (Thermo Fisher Scientific, Waltham, MA) or N-ethylmaleimide (NEM) (Thermo Fisher Scientific, Walthman, MA) for irreversible cysteine thiol binding. Then tardigrades were rinsed twice with 1 mL of deionized water to remove remaining blocking reagent and subjected to centrifugation at 10,000 x*g* for 3 min. Following the second rinse, the supernate was removed, and tardigrades were transferred to 35-mm Petri dishes in less than 20 μL final volume. Tardigrade stressedstress conditions were based on experimentally determined optimized cryptobiosescryptobiosis conditions and times. Post-incubation, tardigrades were assessed for evidence of cryptobiotic state formation as previously described.

### Electron paramagnetic resonance spectroscopy

Aliquots of the superoxide-reactive spin trap 1-hydroxy-3-methoxycarbonyl-2,2,5,5-tetramethylpyrrolidine (CMH, 1 mM final concentration) (Enzo Life Sciences, Farmingdale, NY) were prepared in modified Krebs-HEPES buffer containing 99.01 mM NaCl, 4.69 mM KCl, 2.50 mM CaCl_2_×2H_2_O, 1.20 mM MgSO_4_×7H_2_O, 29.76 mM NaHCO_3_, 1.04 mM KH_2_PO_4_, 11.10 mM D+ glucose, 20.00 mM Na^+^-HEPES, 25 μM deferoxamine mesylate, and 5 μM diethyldithiocarboxamic acid ammonium salt (all reagents from either Thermo Fisher Scientific or Sigma Aldrich). CMH aliquots were stored at -20°C until use. For each of 7 trials (stressed, n=3; non-stressed, n=4), a sample of 200 *H. exemplaris* tardigrades with minimal algae was collected and centrifuged at 13,400 x*g*. The resultant supernate was removed until the final volume of culture sample remaining was 10 μL. For each trial, a stopwatch was started upon transfer of an aliquot of CMH from storage to ice. Prior to adding CMH to samples, the CMH was held under nitrogen gas flow for five minutes to remove dissolved oxygen. Measurements were collected on an X-band EMXPlus Spectrometer (Bruker, Billerica, MA) with the following parameters: sweep width, 100 G; center field, 3510 G; modulation amplitude, 1.000 G; receiver gain, 30; microwave power, 10.02 mW; number of scans, 4. Across trials, samples were time-matched with respect to transferring the CMH to ice (e.g., all samples received CMH at the same time post transfer, were placed into the instrument at the same time post transfer, and data was collected at the same time post transfer). Stressor solution or deionized water and CMH (1:1, v/v; 500 μM working concentration) were added 2 min prior to the collection of data for the initial timepoint. Samples were wicked into 50 μL capillary tubes (Sigma Aldrich, St. Louis, MO). Data points were collected five minutesat 5-min intervals (beginning at 2 min post-addition of CMH/stressor) for 30 minutesmin. Signal intensity for all data points was determined using the difference between the maximum intensity and minimum intensity of the CMH signal and was normalized on a per-animal basis. The rates of CMH oxidation between timepoints were determined using the first derivative of a signal intensity per-animal versus time plot.

### Confocal fluorescent microscopy

Tardigrades (either active or in tuns) were exposed to 30 μM fluorescein-5-maleimide (Sigma Aldrich, St. Louis, MO, USA) in 50 mM PBS, pH 7.2, in 12% (v/v) methanol for 2 h at room temperature. Following incubation, tardigrades were centrifuged at 10,000 x*g* for 5 min, before removing the supernate and rinsing with 1 mL of 50 mM PBS, 12% methanol (v/v), pH 7.2. Brightfield contrast and fluorescence images were acquired on a Zeiss LSM710 confocal laser-scanning microscope using a PlanApo 40X objective lens. Brightfield and fluorescence images were collected using 561-nm and 480-nm lasers, respectively, on Zeiss ZEN software (Car Zeiss, INC. NY, USA).

### Confocal Bright Field Imaging

#### Sample Preparation

All samples were imaged on microscope slides with tardigrades placed in 20 μL total volume droplets within SecureSeal^™^ Imaging Spacers (Thermo Fisher Scientific, Waltham, MA) to prevent crushing of tardigrades. Hydrated tardigrades were dosed with 12.5% (v/v) methanol in water (Thermo Fisher Scientific, Waltham, MA) for control images. Application of osmobiosis and chemobiosis for imaging was identical to as previously described. After animals were placed inside spacer, the slides were sealed with a glass coverslip.

Cryobiote preparation on slides varied. Animals were placed in droplets in minimal media (10 μL) on a 35-mm plastic petri dish and cryobiosis was induced as previously described. 10 μL of thawed but cold 25% (v/v) methanol solution in water previously kept at -80° C was added to the droplet containing bears. Once the entire mixture had thawed, all 20 μL were plated on slides in spacers and sealed.

#### Imaging

All images were obtained using a Leica SP5 TCSII (Wetzlar, Germany). 10 individuals were imaged for each condition. Tardigrade sizes were determined by measuring from the most anterior region to the most posterior region on ImageJ.^34^

### Protein extraction

Tardigrade proteins were extracted using a modified acid extraction.^35^ Tardigrades were incubated in concentrated trifluoroacetic acid (Sigma Aldrich, St. Louis, MO) for 40 min before neutralizing to approximately pH 7 with 2 M Tris (Thermo Fisher Scientific, Waltham, MA) in water. Proteins were precipitated in 70% (v/v) ethanol (Thermo Fisher Scientific) overnight at - 20 °C before centrifuging at 3,220 x*g* for 1 h and collecting the resulting pellet.

### Quantification of reduced thiols

*H. exemplaris* (1000 animals per sample, n=4 per condition) collected in microcentrifuge tubes were induced into osmobiosis by 75 mM CaCl_2_ and 600 mM sucrose and into cryobiosis as previously described. Following cryptobiote induction for each respective condition, proteins were immediately extracted. For those induced into cryobiosis, frozen samples were homogeneized via a plastic pestle before being subjected to protein extraction. Following precipitation, protein pellets were resuspended in 4 M urea in PBS at pH 7.5 and quantified with a CB-X assay (Biosciences) against BSA standards. Proteins were enzymatically digested with 0.5 ug Trypsin Gold (Promega) for 16 h, shaking at 850 RPM at 37°C. Samples were dried in a CentriVap (Labconco) and subsequently resuspended in PBS at pH 7.5. Resuspended samples were subjected to reduced thiol quantitation using the Measure-iT^™^ Thiol Assay Kit (ThermoFisher) against reduced glutathione following the manufacturer’s protocol. Briefly, the Measure-iT^™^ thiol quantitation reagent was diluted 1:100 in PBS. In a 96 well plate, 100 uL of the diluted thiol quantitation reagent were added to 10 uL of sample/standard. Samples were incubated at RT for 5 min before fluorescence was measured using a SpectraMax M5 (Molecular Devices) microplate reader with excitation/emission maxima at 490/520 nm. Thiol concentrations were normalized to total protein amounts in each sample. Statistical testing was performed using a Student’s t-test.

### Chemical inhibition of tardigrade tuns

Tardigrades were plated one organism per well in a clear 96-well plate in 1-uL droplets of water. Tardigrades were incubated for 1 h with a working concentration of 100 uM of all inhibitors from the SCREENWELL Redox Library (Enzo Life Sciences, Farmingdale, NY) and monitored for physiological changes. Tardigrades were incubated in a working concentration of 75 mM CaCl_2_ and observed for tun formation after 3 h.

Glutathione (reduced and oxidized), L-ergothioniene, and B-lapachone were studied using 30 tardigrades per sample in 4 replicates per inhibitor. Each was applied with a working concentration of 75 mM and 0.1% DMSO to improve permeation. Following incubation for 1 h, tardigrades were treated in 75 mM CaCl_2_ and observed for tun formation.

Erastin inhibition was conducted using 3 replicates of 30 tardigrades, each in an individual well of a 96 well plate. Tardigrades were incubated in a 100 nM working concentration of erastin in 0.1% DMSO overnight. After 20 h, tardigrades were monitored for the formation of tuns.

## Results and Discussion

### ROS exposure induces tun formation in tardigrades–direct evidence of chemobiosis

Tardigrades survive multiple stressors that are prone to generating ROS *in vivo*, including desiccation, extreme fluctuation in osmolarity, and freezing, among others.^14, 36-39^ We therefore hypothesized that ROS may play a crucial role in signaling tun induction in response to environmental fluctuations. This was explored using exogenously applied H_2_O_2_ and monitoring the physiological effect(s) on tardigrade morphology (**Figure 1**). Tardigrades exhibited a dose-dependent response to exogenous H_2_O_2_, with higher concentrations generating tuns within an of exposure. Tardigrades exposed to peroxide retracted their limbs and decreased their size to approximately 25% of their hydrated counterparts, thus indicating chemobiosis resulting in tun formation had taken place (**Figure 1A**). To further explore this phenomenon, tardigrade tun formation was monitored over time for varying concentrations. Measurements were conducted in 3 biological replicates, with a single replicate including 25-30 specimens. Time to tun formation was distinct across peroxide concentrations, with 5 mM H_2_O_2_ resulting in tuns for 10% of the tardigrades by 90 min of exposure, while 750 μM H_2_O_2_ took at least 4 h to achieve the same result (**Figure 1B**). All tardigrades entered tuns following exposure to 750 μM H_2_O_2_ by 12 h, whereas it took only 8 h with 2.5 mM H_2_O_2_. Only 57% of tardigrades exposed to 5 mM H_2_O_2_ entered tuns; the remaining 43% entered a turgid, semi-translucent state indicative of tardigrade death. The lethality of higher ROS concentrations could indicate a risk of over-oxidation (perhaps in the form of irreversible oxidation, in contrast to reversible) in extreme conditions. This was further mirrored by a significantly decreased tardigrade viability at all concentrations >1 mM H_2_O_2_, with only 34% survival following exposure to 2.5 mM H_2_O_2_ and 16% following exposure to 5 mM H_2_O_2_, both determined by recovery of motility following emergence from tuns (**Figure 1C**). In contrast, concentrations of 1 mM and 750 μM H_2_O_2_ resulted in post-tun tardigrade viability of 67.4 and 70.0%, respectively.

**Figure 1.**
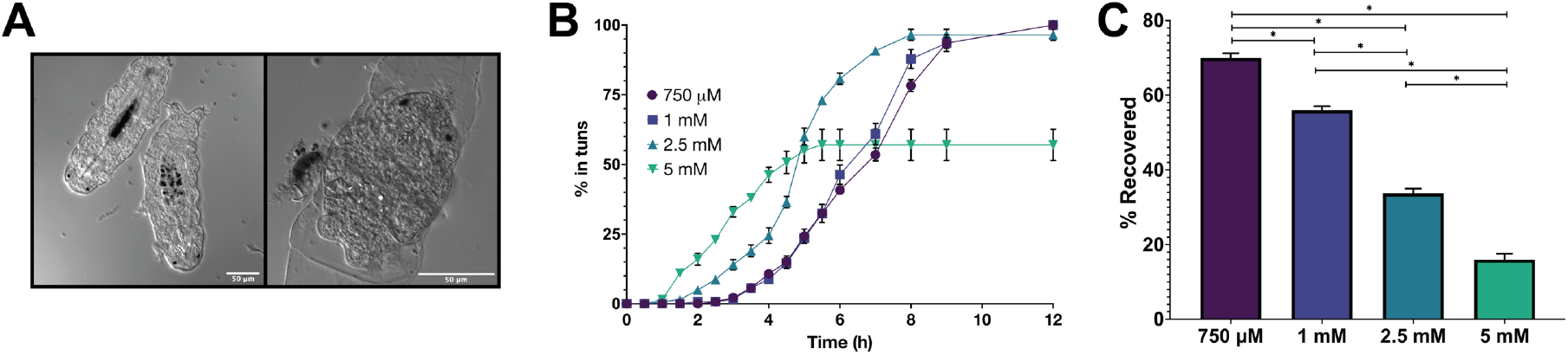
Peroxide induces tuns in a concentration dependent manner in *Hypsibius exemplaris*. A) Confocal images of tardigrades in the hydrated state as well as a H_2_O_2_– induced tun. Tuns are about 25% of the size of the hydrated tardigrades. B) The percent of tardigrades entering tuns following different concentrations of H_2_O_2_. Tardigrades were measured in three biological replicates, where a single replicate included 25 – 30 tardigrades. Error bars represent standard deviation. C) The percentage of tardigrades that reemerged from H_2_O_2_– induced tuns following reintroduction to distilled water. Error bars represent standard deviation. Only tardigrades that enter tuns are counted toward total survival. Statistical significance was determined by paired t-tests with Welch’s correction, where ^*^ indicates p < 0.05.

Despite the absence of desiccation or significant fluctuations in osmolarity, tardigrades still expelled internal water stores and initiated tun formation, resulting in an approximate 75% decrease in body length (**Figure 1A**). This suggests that water expulsion during tun formation is an active process rather than the result of spontaneous flux, and that the mechanism(s) of expulsion could be mediated by oxidation or oxidative signaling. This process was quickly reversed when tuns were placed back in distilled water in the absence of exogenous ROS, suggesting that tardigrades are able to rapidly sense environmental changes to initiate anatomical transitions. Tardigrades’ ability to undergo chemobiosis has previously been predicted, but the data presented here is the first empirical evidence of chemical-induced tun induction.

### *H. exemplaris* can be reproducibly induced into cryo- and osmobiosis

While cryobiosis has been previously explored in *H. exemplaris*, osmobiosis required thorough empirical characterization. We mapped the induction and concentration of osmobiosis using two distinct osmolytes, CaCl_2_ and sucrose, chosen due to their ionic and non-ionic natures (**Figure 2**). Both stressors were applied in concentrations far exceeding expected physiological concentrations, as it was essential to induce a stress response that would prove fatal should the tardigrades not enter their resilient tun form. Furthermore, tardigrades do not form tuns in response to physiological conditions; rather, they must be exposed to extreme stress to initiate their tun formation. As observed with peroxide, tun inductions by CaCl_2_ and sucrose were highly concentration dependent. While 50 mM CaCl_2_ resulted in total tun formation in 12 h, 75 mM CaCl_2_ produced the same result in only 3.5 h (**Figure 2A**). Exposure to 100 and 150 mM CaCl_2_ resulted in 92% and 52% of tuns induced after 1 h, respectively; interestingly, these higher concentrations also resulted in tardigrade fatality before tun formation could occur. Survival was high in all concentrations for those that formed tuns following CaCl_2_ exposure, with an average survival rate of 80 – 90% for all concentrations (**Figure 2C**). Sucrose also formed tuns in a concentration-dependent manner (**Figure 2B**); however, unlike peroxide and CaCl_2_ exposure, higher concentrations did not result in tardigrade death prior to tun formation. This is consistent with osmobiosis mapped in the tardigrade species *Ramazzottius oberhaeuseri*, which had a higher tolerance to nonionic osmolytes in comparison to ionic.^39^ All tardigrades exposed to 225 mM, 300 mM, 450 mM, and 600 mM sucrose formed tuns, with 100% of the tardigrades forming tuns at 5.5, 4, 2, and 0.5 h, respectively. Interestingly, the survival rate post-tun formation was not concentration dependent, with repeated trials producing the same pattern of survival that is inconsistent with dose concentration (**Figure 2D**). While deviation in tardigrade size and/or age could generate stochasticity in the data, these differences would be expected to be random. The repeated findings of non-correlative survival rates suggest that this is a regulated phenomenon rather than random.

**Figure 2.**
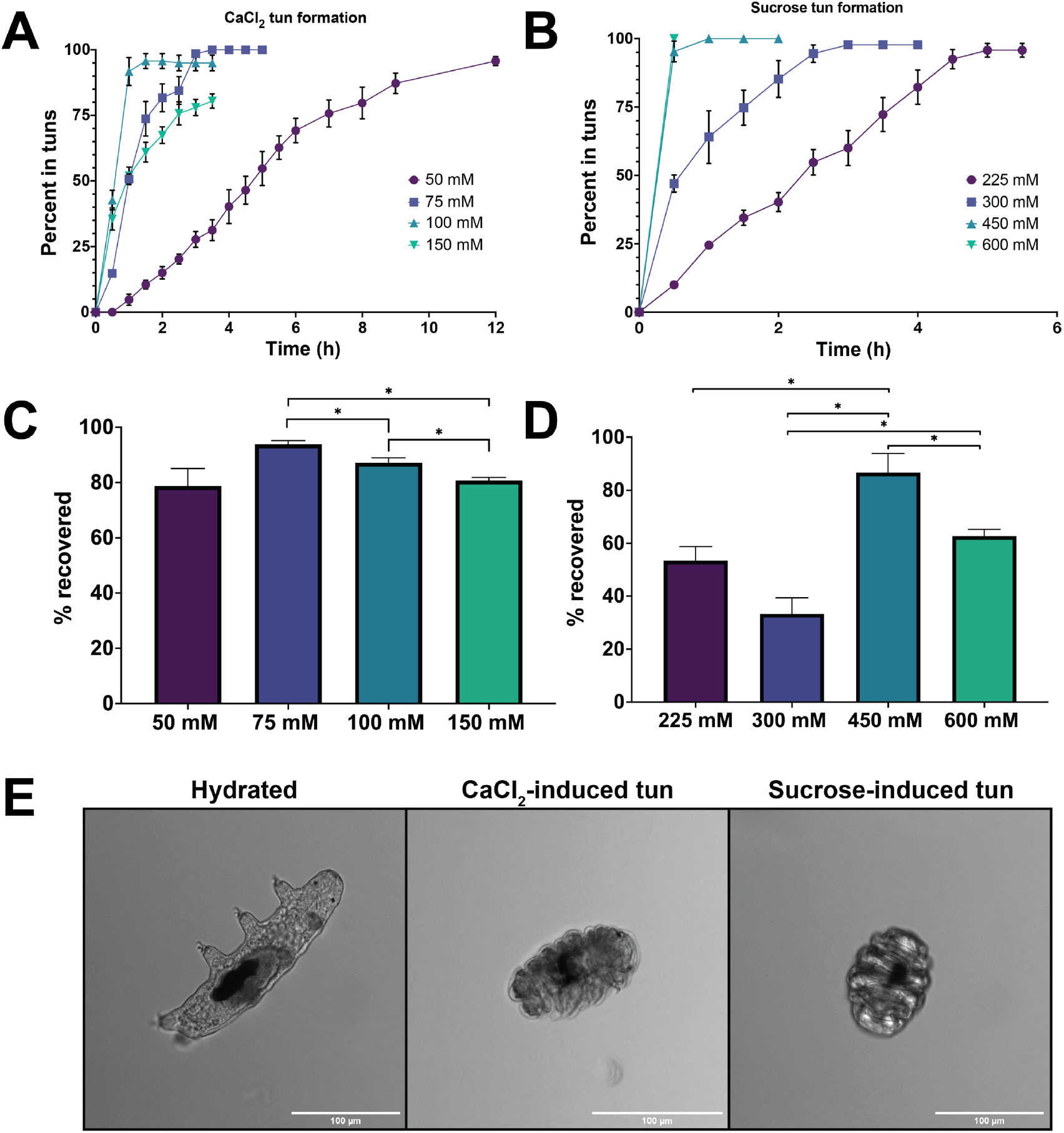
Tardigrades can be induced via osmobiosis using both CaCl_2_ and sucrose. A) The percent of tardigrades entering tuns following different concentrations of CaCl_2_. Tardigrades were measured in three biological replicates, where a single replicate included 25 – 30 tardigrades. B) The percent of tardigrades entering tuns following different concentrations of sucrose. Tardigrades were measured in three biological replicates, where a single replicate included 25 – 30 tardigrades. C) The percentage of tardigrades that reemerged from CaCl_2_–induced tuns following reintroduction to aqueous media. D) The percentage of tardigrades that reemerged from sucrose–induced tuns following reintroduction to aqueous media. A-D) Error bars represent standard deviation. C-D) Statistical significance was determined by paired t-tests with Welch’s correction, where ^*^ indicates p < 0.05. E) Tardigrades imaged using a confocal microscope in their hydrated, CaCl_2_-induced, and sucrose-induced states.

Both CaCl_2_ and sucrose can reproducibly produce tuns with high rates of post-stressor survival (**Figure 2**). Physiologically, there appears to be slight differences between CaCl_2_ and sucrose-induced tuns, with sucrose tuns appearing more compact (**Figure 2E**). This could be due to the higher calculated osmotic pressure (π) in sucrose, with 15.17 bar compared to 9.59 bar; however, more investigation is needed to determine the precise cause. Interestingly, this tun formation and survival was not consistent across all osmolytes. Glucose and glycerol exhibited nearly total tun formation at 600 mM, but survival was less than 50%; in contrast, neither Na Cl or MnCl_2_ formed reproducible tuns in repeated trials (data not shown). This suggests that osmotic pressure alone is not the inducer of tun formation, and that intracellular regulatory mechanisms must have some control over tun initiation.

### Reversible cysteine oxidation is essential for tun formation

ROS contribute to cellular homeostasis through rapid reversible oxidative signaling through which the oxidation of cysteine thiols is used to trigger protein responses.^40^ Therefore, ROS induction of tun formation through the oxidation of cysteine thiols was examined by blocking reduced cysteine thiols in hydrated tardigrades with either iodoacetamide (IAM) or *N*-ethylmaleimide (NEM), both of which irreversibly bind to reduced cysteine thiols and block oxidation.^41^ Tardigrades were incubated in either 30 μM IAM or NEM (concentration chosen following titration series to ensure tardigrade survival) and exposed to 750 μM H_2_O_2_ for 12 h (**Figure 3A**). Experiment was conducted with 3 biological replicates, where each biological replicate included 20 specimens. While 100% of tardigrades formed tuns in the absence of Cys-blocking, only 7.6 and 4.0% of tardigrades blocked with IAM and NEM, respectively, formed tuns by 12 h. This dramatic decrease in tun formation indicates an essential role for cysteine oxidation in the mechanism and/or signaling of the process. While a small difference was observed in IAM compared to NEM, this difference was not statistically significant, and likely represents the slightly higher binding affinity and specificity of NEM compared to IAM.^42, 43^ Tardigrades exposed to NEM before being exposed to peroxide showed decreased size compared to fully hydrated, and visible deformity with limbs extended rather than retracted (**Figure 3B**), suggesting that cysteine oxidation is necessary for limb retraction during tun formation.

**Figure 3.**
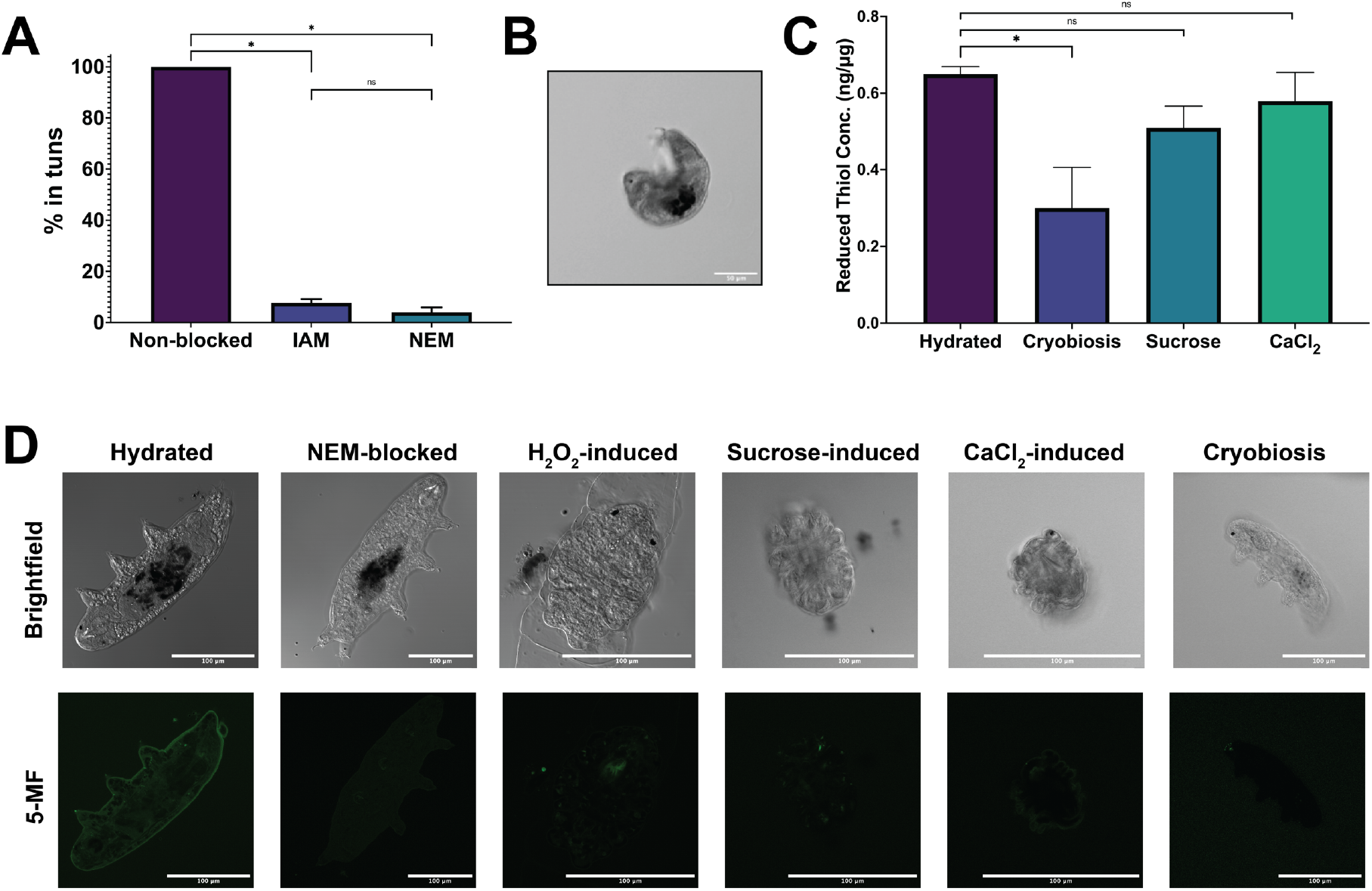
Blocking of cysteine residues prevents tun formation. A) Tardigrades exposed to either IAM or NEM, both of which bind irreversibly to reduced cysteines, significantly decreased tun formation in H_2_O_2_–exposed tardigrades. B) Confocal image of a NEM-blocked tardigrade following exposure to H_2_O_2_. C) Tardigrade proteins were extracted and digested with trypsin before fluorescent quantification of reduced cysteine thiols. Thiol concentration was normalized to protein concentration. A, C) Error bars represent standard deviation. Statistical significance was determined by paired t-tests with Welch’s correction, where ^*^ indicates p < 0.05. D)Tardigrades were labeled with 5-MF before imaging with a confocal microscope. To demonstrate the selectivity of the label, one set of tardigrades was blocked with NEM prior to 5-MF labeling, showing significantly diminished fluorescence compared to the unblocked tardigrade.

Whereas chemo- and osmobiosis forms tuns, cryobiosis does not; it was therefore unclear whether or not oxidation would still be implicated in cryobiosis. To assess this phenomenon, cryobiosis was analyzed to ensure high survival rates, with a freezing rate of 1°C/min resulting in 87% survival (**Supplemental Figure 1A**). Cryobiotes imaged using a confocal microscope revealed a distinct morphology from osmo- or chemobiotes, with the tardigrades appearing more globular rather than the ordered, compact form of tuns (**Supplemental Figure 1B**). To determine if cysteine oxidation was necessary for survival, tardigrades were exposed to 30 uM NEM prior to freezing. This resulted in tardigrades becoming more linear and distended during freezing, likely contributing to the lack of any survivors following reheating to room temperature (**Supplemental Figure 1C**). Based on this data, it appears that cysteine oxidation is still necessary for cryptobiotic survival, even when the tun formation is not involved.

In addition to blocking reversible oxidation of cysteine thiols using NEM or IAM, cryptobiosis can be blocked using 12.5% (v/v) methanol. Tun formation was inhibited by 91% and 99% in sucrose- and CaCl_2_-exposed samples, respectively, and cryobiote formation was inhibited by 100% (**Table 1**). The blocking of tun formation by methanol gives further evidence that tun initiation is not the result of desiccation via passive osmosis; rather, it is a highly regulated phenomenon.

**Table 1.**
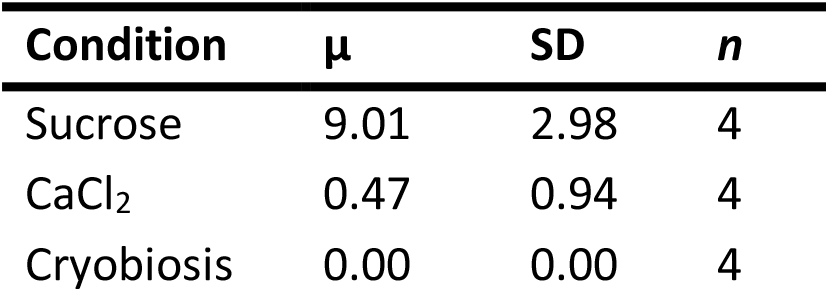
Percentage of tardigrades that undergo either osmobiosis or cryobiosis following exposure to methanol. Biological replicates included 30 specimens each. None of the cryobiotic tardigrades survived.

| Condition | $\mu$ | SD | *n* |
| --- | --- | --- | --- |
| Sucrose | 9.01 | 2.98 | 4 |
| CaCl<sub>2</sub> | 0.47 | 0.94 | 4 |
| Cryobiosis | 0.00 | 0.00 | 4 |

For quantitative assessment of thiol oxidation, reduced thiols were quantified from the tardigrade proteome was quantified using a fluorescence-based quantitative thiol assay. Proteome digestion was essential to increase accessibility to internal cysteines that may be sterically hindered from binding to the fluorophore. To circumvent variation due to tardigrade size and/or age, quantified cysteine thiols were normalized to quantified protein content. All cryptobioses exhibited lower concentrations concentration of reduced cysteines (**Figure 3C**). Cryobiosis had the lowest of all stressors, with approximately half of the free cysteines compared to the control (**Figure 3C**). This suggests differential cysteine regulation across stressors.

Further qualitative assessment of cysteine thiol oxidation was conducted using confocal microscopy with fluorescein-5-maleimide (5-MF), a labeling agent that irreversibly derivatizes cysteine thiols, across all inducible stressors (**Figure 3D**). Active tardigrades exhibited fluorescence, with 5-MF localized near the tardigrade epidermis, whereas tuns and cryobiotes exhibited minimal fluorescence. This indicates that reduced cysteine thiols in the active, hydrated tardigrades were likely oxidized and inaccessible to 5-MF labeling once in the tun state. As with the thiol quantification assay, cryobiosis exhibited the lowest fluorescence with 5-MF. Interestingly, the remaining cryptobiotes also have diminished fluorescence compared to the control.

Cysteines can rapidly undergo a variety of reversible oxidative modifications, including sulphenylation, disulfide bond formation, glutathionylation, and *S-*nitrosylation, that often facilitate rapid intracellular signaling.^44, 45^ To determine the reversibility of tun-inducing cysteine oxidation, tuns induced by 12-h exposure to 750 μM H_2_O_2_ were exposed to the reductant β-mercaptoethanol (BME). After 30 min of concentrated (14.3 M) BME exposure, 98% of tardigrades had reemerged to their linear state, with emergence visualized via microscopy beginning at 5 min. Tuns induced by up to 500 mM H_2_O_2_ similarly reemerged from tuns into a linear state. Interestingly, reductant-induced tun release was lethal to all tardigrades, indicating that native tun release must be a highly regulated physiological event. This could potentially occur through crosstalk with other post-translational modifications; the rapid recovery of H_2_O_2_-induced tuns following placement in distilled water (<1 h) likely precludes an upregulation in protein expression as a contributing factor. The reversibility of tuns using BME suggests that the cysteine modification required for tun formation is reversible oxidation that can be controlled through exogenous ROS and reducing agents.

### Tardigrades may use an intricate network of ROS signals to initiate and survive tun formation

Our results indicate that the reversible oxidation of cysteine thiols is essential for tun induction in *H. exemplaris*. The dose-dependence of tun formation in the presence of H_2_O_2_ suggests that either environmental ROS or the intracellular production of ROS in the presence of external stressors triggers tardigrades to enter cryptobiosis. To further assess intracellular ROS generation in tun formation, tardigrades were screened using the SCREEN-WELL REDOX library, through which they were exposed to either antioxidants or oxidants, monitored for physiological changes, and then induced into tuns using CaCl_2_. Compounds that did not have a physiological effect alone but prevented tun formation following stressor introduction were examined further. In total, 84 compounds were screened, including many phenolic antioxidants, radical scavengers, and thiol-containing reducing agents. Following application of CaCl_2_, 72 of the compounds inhibited tun formation, with 15 inhibiting over 50% of tardigrades (**Supplementary table 1**). Three of the highest inhibiting compounds in this preliminary screen were selected for re-analysis across replicates: glutathione, L-ergothioneine, and β -lapachone.

Glutathione was one of the largest inhibitors of tun formation, preventing tuns in ∼68% of the tardigrades following exposure to CaCl_2_ (**Figure 4**). When oxidized glutathione was used instead, tun inhibition decreased to 22%, thus indicating that reduced glutathione has a significant effect on tun formation. This likely derives from the readily oxidized cysteine in glutathione where an influx of glutathione renders ROS unable to reach their target(s). Two other antioxidants were evaluated for their ability to prevent tun formation: L-ergothioneine, a sulfur containing small molecule, and β -lapachone, a sulfur-free benzochromenone, both of which significantly inhibited tun formation, with 52% and 69% inhibition, respectively (**Figure 4B**). The ability to suppress tun formation with antioxidants provides evidence that intracellularly generated ROS mediate tardigrade survival in the presence of exogenous stressors.

**Figure 4.**
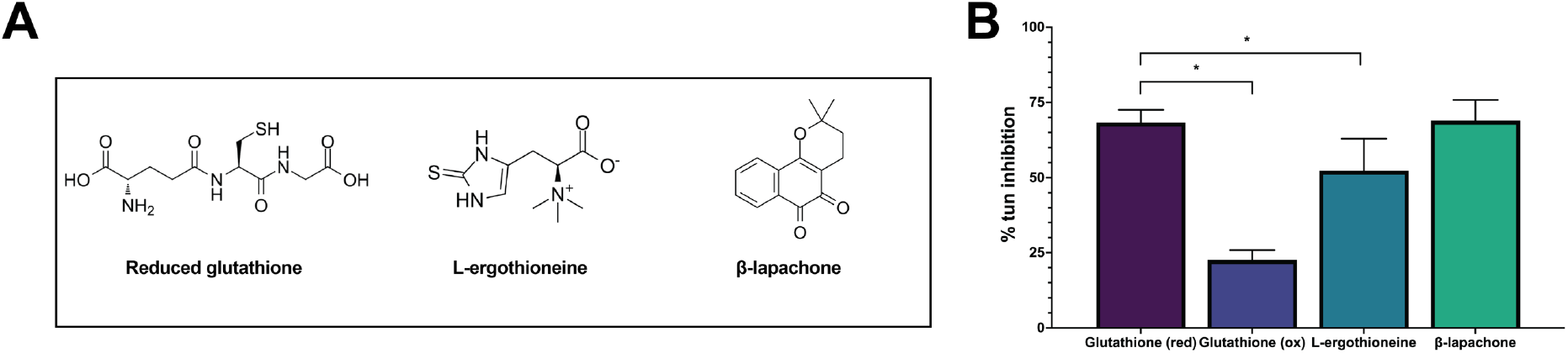
Exposure to exogenous antioxidants prior to osmotic stress decreases tardigrade survival. A) Structures of antioxidants. B) Three inhibitors were further characterized in replicates following the initial screening. Glutathione, L-ergothioneine, and B-lapachone all inhibited >50% of tuns from forming. Error bars indicate standard deviation. Statistical significance was determined by paired t-tests with Welch’s correction, where ^*^ indicates p < 0.05.

EPR spectroscopy was implemented to further delineate ROS accumulation following exposure to stress. Hydrated, hydrated tardigrades and those exposed to 600 mM sucrose were analyzed using the superoxide-sensitive spin probe CMH.^46^ Superoxide is generated by electron leak at Complex I and Complex III of the mitochondria, the rate of which is increased under stress.^47, 48^ CMH is a hydroxylamine probe that generates a stable nitroxide radical that can be detected via EPR (**Figure 5A**). Exposing CMH to hydrated and sucrose-exposed tardigrades thus enabled relative quantification of superoxide production over time, allowing us to compare the rate of accumulation as tardigrades entered their osmobiotic tun. Both hydrated and sucrose-exposed tardigrades started with overlapping rate of superoxide accumulation (**Figure 5B**). By 10 min, however, sucrose significantly increased in accumulation rate compared to the hydrated sample, with an approximately 25% increase in superoxide rate. This increase continued for the duration of the 30-min measurements, thus showing that the induction of osmobiosis releases intracellular ROS.

**Figure 5.**
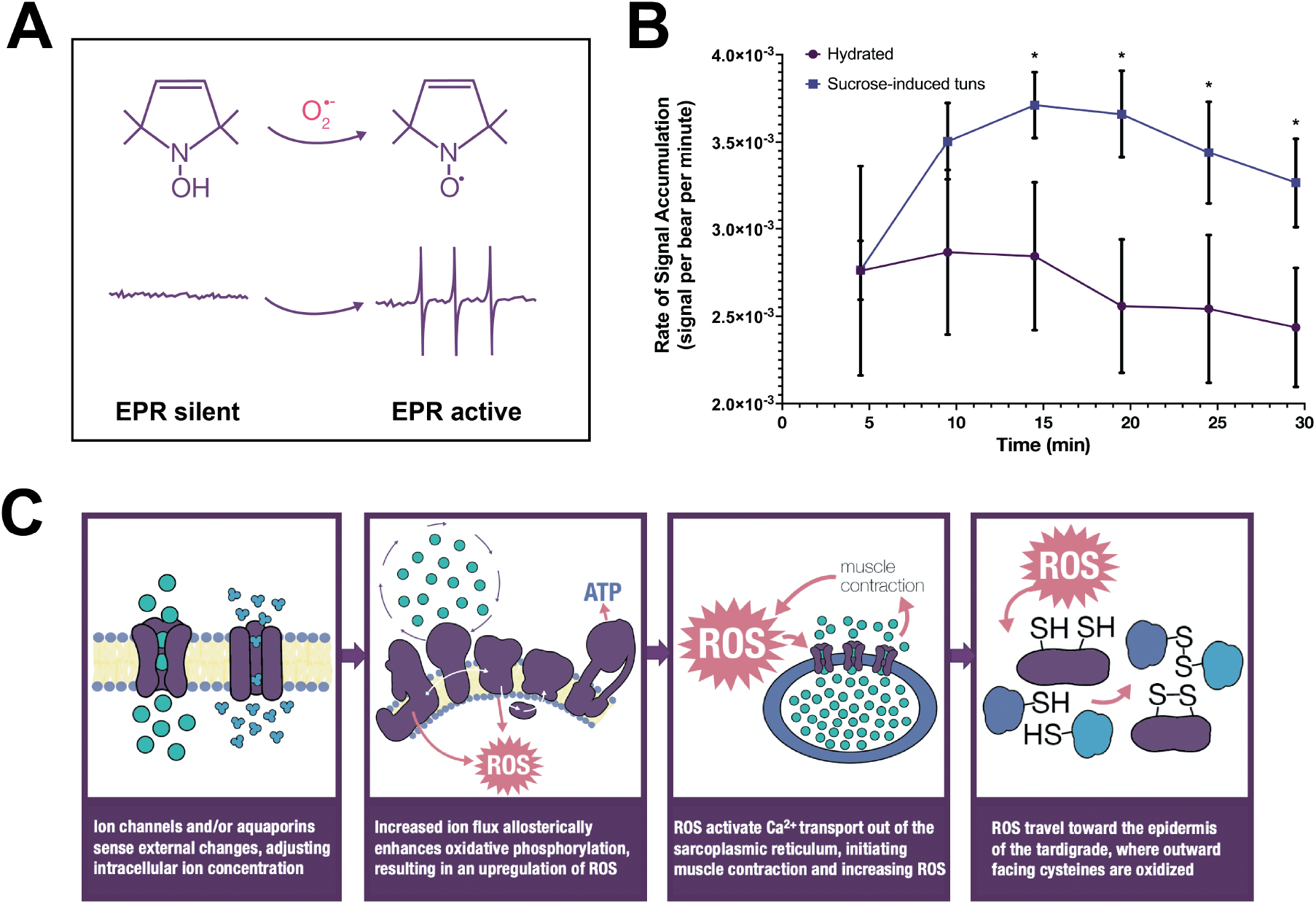
Tardigrades accumulate ROS following stress exposure. A) CMH forms a stable nitroxide radical following contact with superoxide that is quantifiable via EPR. B) Hydrated and sucrose exposed tardigrades were probed with CMH and measured with EPR every 5 min. Sucrose exposed tardigrades accumulate superoxide at significantly higher rates than non-stressed tardigrades, beginning at 10 min post stress introduction. Error bars indicate standard deviation. Statistical significance was determined by paired t-tests at each time point with Welch’s correction, where ^*^ indicates p < 0.05. C) The working hypothesis for tardigrade survival via ROS-mediated protein signaling.

Mitochondrial ROS are modulated by ion fluxes (particularly calcium), which allosterically increase mitochondrial electron transport via complex II (**Figure 5C**). We therefore hypothesized that a critical regulatory protein in tardigrade survival would be voltage-gated anion channel 2 (VDAC2), a highly abundant mitochondrial channel responsible for the majority of calcium flux in and out of the mitochondria. To determine the influence of VDAC2 on tun formation and tardigrade survival, tardigrades were exposed to the VDAC2 inhibitor erastin and monitored for physiological changes. Following exposure to 100 nM of erastin, tardigrades began to form tuns within minutes. Within 24 h, 75.56% (SD: 6.94) of the tardigrades had formed tuns. This suggests that the flux of calcium in and out of the mitochondria is likely implicated in tun formation and that VDAC2 closes to decrease flux in response to external stressors.

## Conclusion

Tardigrades have long been renowned for their remarkable ability to survive adverse conditions, an achievement that has helped them to survive fluctuating environments since they evolved in the Cambrian era. However, the mechanisms used to sense and respond to external stressors in order to initiate survival strategies have not previously been described. We have revealed that tardigrade survival is dependent on reversibly oxidized cysteines coordinating the entrance and emergence from survival states in a highly regulated manner. Through implementation of EPR and redox library screens, we have demonstrated that intracellular release of ROS is essential for tun formation. We have also characterized the initiation, emergence from, and dose-dependence of both osmobiosis and chemobiosis, shown here in *H. exemplaris* for the first time. The rapid induction of tardigrade tuns via osmobiosis, paired with the high survival rates following emergence, makes it an easily implemented cryptobiotic state for further exploration of the mechanisms of tardigrade survival.

## Acknowledgements

This research was supported by National Science Foundation grants awarded to L.M.H. (NSF-MCB 2149172) and D.R.J.K. (NSF-MCB 2149173). A.L.S. acknowledges funding from the North Carolina Space Grant. Marshall University students were also funded by a National Science Foundation (NSF) Grant (Award Nos. CHE1229498 and OIA1458952), the NASA West Virginia Space Grant Consortium (Grant no. NNX15AK74A).

## Author Contributions

**Conceptualization**: ALS, DRJK, JRC, LMH; **Methodology**: ALS, BBF, DRJK, ERS, HMO, JRC, KMJ, LMH, TAC; **Validation**: BBF, HMO, KMJ, MHD, TAC; **Formal Analysis**: ALS, BBF, ERS, HMO, JRC, KMJ; **Investigation**: ALS, BBF, ERS, HCL, HMO, JRC, KMJ, MHD, SNM, TAC; **Resources**: BBF, DRJK, HMO, KMJ, TAC; **Data Curation**: ALS, DRJK, KMJ, LMH; **Writing-Original Draft**: ALS, KMJ; **Writing – Review & Editing**: ALS, BBF, DRJK, KMJ, LMH; **Visualization**: ALS; **Supervision**: DRJK, LMH; **Project Administration**: DRJK, LMH; **Funding Acquisition**: DRJK, LMH.

## Conflict of Interest

The authors declare no conflicts of interest.

## Notes

### Competing Interest Statement

The authors have declared no competing interest.

