## Supplemental Figure 1 for "Chemobiosis reveals tardigrade tun formation is dependent on reversible cysteine oxidation"

A

| *n* | Survival % |
|---|---|
| 30 | 87% |
| 30 | 80% |
| 31 | 90% |
| 30 | 87% |
| 32 | 94% |

$\mu = 87.6 \pm 5.1\%$

B C

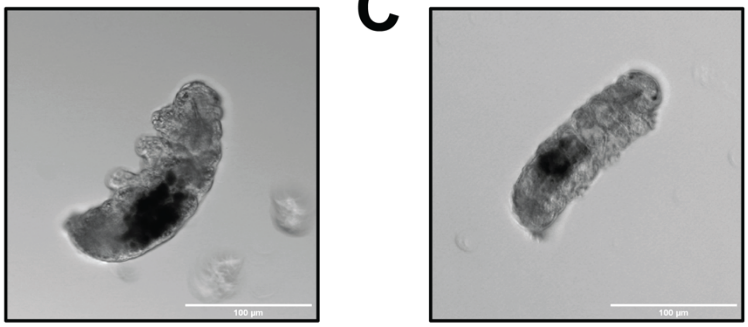

**Supplemental Figure 1.** Tardigrades rely on cysteine oxidation to survive cryobiosis. **A**) The survival of tardigrades that underwent cryobiosis for 24 h. Survival was determined by coordinated leg movements following return to room temperature. **B**) Confocal image of a tardigrade undergoing cryobiosis. Note that cryobiosis does not facilitate tun induction. **C**) Confocal image of a tardigrade exposed to 30 uM NEM prior to undergoing cryobiosis. None of the NEM-blocked tardigardes survived cryobiosis.
